# SLIMEr: probing flexibility of lipid metabolism in yeast with an improved constraint-based modeling framework

**DOI:** 10.1101/324863

**Authors:** Benjamín J. Sánchez, Feiran Li, Eduard J. Kerkhoven, Jens Nielsen

## Abstract

A recurrent problem in genome-scale metabolic models (GEMs) is to correctly represent lipids as biomass requirements, due to the numerous of possible combinations of individual lipid species and the corresponding lack of fully detailed data. In this study we present SLIMEr, a formalism for correctly representing lipid requirements in GEMs using commonly available experimental data. SLIMEr enhances a GEM with mathematical constructs where we Split Lipids Into Measurable Entities (SLIME reactions), in addition to constraints on both the lipid classes and the acyl chain distribution. By implementing SLIMEr on the consensus GEM of *Saccharomyces cerevisiae*, we can predict accurate amounts of lipid species, analyze the flexibility of the resulting distribution, and compute the energy costs of moving from one metabolic state to another. The approach shows potential for better understanding lipid metabolism in yeast under different conditions. SLIMEr is freely available at https://github.com/SysBioChalmers/SLIMEr.

## 1. Background

Genome scale metabolic models (GEMs) are widely used to model and compute functional states of cellular metabolism (Nielsen, 2017). A crucial step for achieving these simulations is to define a biomass pseudo-reaction (Feist & Palsson, 2010; Dikicioglu *et al*, 2015), which accounts for every single constituent comprising the cellular biomass (proteins, carbohydrates, lipids, etc.). In this step it is challenging to account for lipid requirements, as there are copious different individual lipid species: over 20 different classes of lipids can be produced in a cell, and each specific lipid belonging to any of those classes can contain various combinations of acyl chain groups, each of them with varying length and number of saturations (Ejsing *et al*, 2009). This can yield over 1000 specific lipid species that the cell can potentially produce. Unsurprisingly, lipid metabolism therefore tends to be the most complicated part of any GEM.

A necessary requirement for formulating the biomass pseudo-reaction is to have abundance measurements [mmol/gDW] of every single constituent; however, this is seldom available for individual lipid species; what is more common is to instead measure separately (i) a profile of all different lipid classes, with high-performance liquid chromatography (Khoomrung *et al*, 2013), and (ii) a distribution of all different acyl chains, with fatty acid methyl ester (FAME) analysis (Abdulkadir & Tsuchiya, 2008). GEMs need to therefore handle these types of data.

The most common approach to represent lipid metabolism in GEMs is to enforce a specific distribution of each individual lipid species, either by using detailed experimental data (Mardinoglu *et al*, 2014; Lachance *et al*, 2018) or by assuming that lipid classes have all the same acyl-chain distribution from a single FAME analysis (Nookaew *et al*, 2008; Kerkhoven *et al*, 2016). In both cases however, the model will be fixed to follow a predefined lipid distribution. This is undesirable, as lipid metabolism can show a high level of reorganization (Ejsing *et al*, 2009; Han *et al*, 2011), hence rendering the model’s predictions of limited use when simulating different experimental conditions, or when looking into the network’s flexibility for satisfying lipid requirements.

A second popular approach is to allow any specific lipid to form a corresponding generic lipid class (e.g., “phosphocholine”) and to only constrain those classes with experimental abundances from lipid profiling (Heavner *et al*, 2012; Brunk *et al*, 2018). The problem with this approach is that experimental abundances from FAME analysis are neglected, and simulations always end up choosing lipid species that cost the least energy, which might not reflect reality, e.g. if there is regulation in place to ensure production of longer chain species. Hence, there is need for an approach that can incorporate both lipid profiling and FAME analysis, but at the same time can allow flexibility in the metabolic network.

In this work, we introduce SLIMEr, a method for correctly representing lipid requirements in GEMs while allowing network flexibility. The approach adds so-called SLIME reactions, which split lipids into their basic components; and lipid pseudo-reactions, that impose constraints on both the lipid classes and the acyl chain distributions. By following this approach, we achieve flux simulations that respect both the lipid class and acyl chain experimental distribution, and at the same time avoid over-constraining the model to only simulate one lipid distribution. We implemented this approach for the consensus GEM of *Saccharomyces cerevisiae* (budding yeast), a model that has undergone iterative improvements (Herrgård *et al*, 2008; Dobson *et al*, 2010; Heavner *et al*, 2012, 2013; Aung *et al*, 2013) and is currently being hosted at https://github.com/SysBioChalmers/yeast-GEM. We show that the enhanced model can better represent lipid requirements for metabolism, has a high degree of flexibility, and can simulate how lipid costs change at varying experimental conditions.

## 2. Methods

### 2.1. FBA and the challenge of representing lipid composition

Flux balance analysis (FBA) (Orth *et al*, 2010) is based on the following assumptions on metabolism: (i) a cell has a metabolic goal, which we can represent through a mathematical objective function, (ii) under short timescales there is no accumulation of intracellular metabolites, and (iii) metabolic fluxes are bounded to physical constraints, such as thermodynamics and kinetics. Those 3 assumptions define a basic FBA problem as followed:

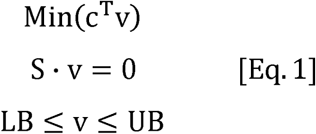

where S is the stoichiometric matrix, which contains the stoichiometric coefficients for all reactions and metabolites, v is the vector of metabolic fluxes [mmol/gDWh], c is the objective function vector, and LB and UB are the corresponding lower and upper bounds for each of the fluxes (some of them based on experimental values). As we usually wish to simulate growth, a biomass pseudo-reaction is typically added to the stoichiometric matrix as follows:

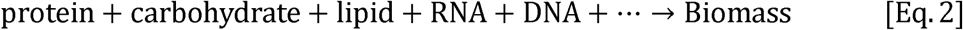

where protein, carbohydrate, lipid, etc., are pseudo-metabolites that are produced from a combination of metabolic components. For example, protein is produced from a protein pseudo-reaction:

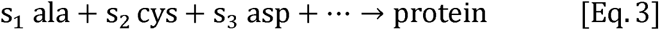

where s_i_ are the measured abundances [mmol/gDW] of the corresponding amino acids in yeast. In this study we focus on the lipid pseudo-reaction, which becomes more challenging to formulate, because there are so many different individual species. One option is to define an equivalent reaction to Eq.3 with every single lipid species (Mardinoglu *et al*, 2014); however, as these measurements are not available for most organisms, the lipid pseudo-reaction is usually represented as the following instead:

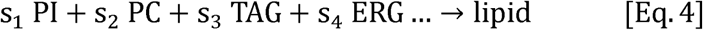

where PI (phosphoinositol), PC (phosphocholine), TAG (triglyceride), ergosterol (ERG), etc. represent each of the lipid classes that exist in the model. Most of them represent not one but a plurality of different molecules, each with different combinations of acyl chain lengths and saturations. Therefore, they are also pseudo-metabolites that need to be produced in turn by additional pseudo-reactions. These pseudo-reactions can be constructed either in a restrictive or permissive approach (Figure 1). The restrictive approach is to enforce the experimental FAME distribution to every single specific lipid species. This can be achieved by creating a generic acyl chain component (Nookaew *et al*, 2008):

**Figure 1:**
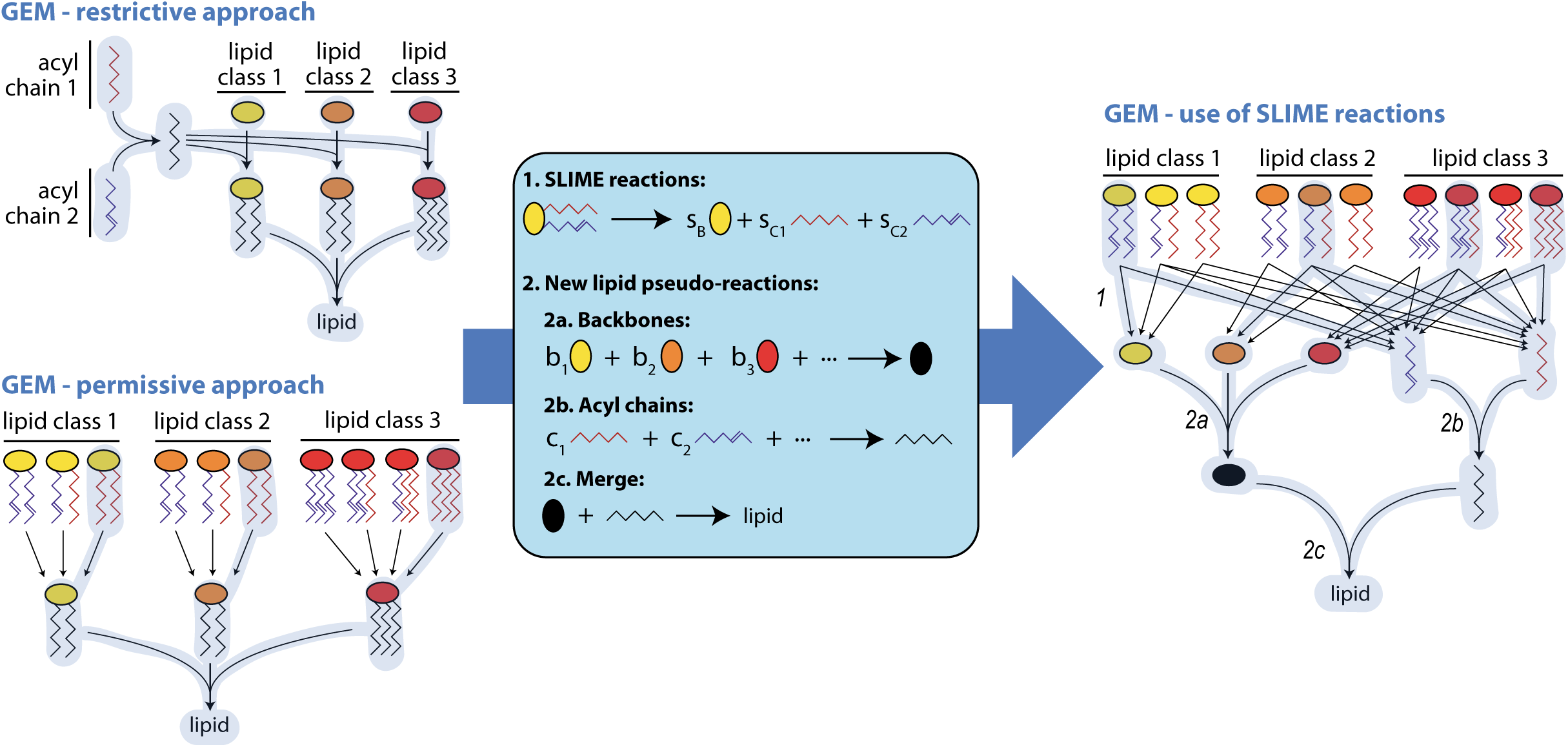
Overview of the process of including SLIME reactions and new lipid pseudo-reactions for a hypothetical model of 3 lipid classes and 2 types of acyl-chain. The active fluxes after simulating the models are highlighted in light blue, showing that a GEM with a restrictive approach would use the same acyl chain composition for all lipid classes (left upper corner), a GEM with a permissive approach would always choose the cheapest species from each lipid class (left lower corner), and a GEM with SLIME reactions would satisfy both the lipid class and the acyl chain distribution, but choosing freely which specific lipid species to produce for this goal (right side).

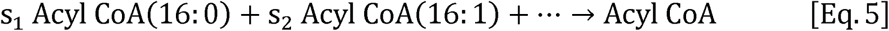

where the s_i_ coefficients are fractions inferred from FAME data. This generic Acyl – CoA is then used to form each generic lipid species, forcing then every lipid class to have the same acyl chain distribution. The latter is an important limitation of this approach, considering that the acyl chain distribution can vary significantly across lipid classes (Ejsing *et al*, 2009).

On the other hand, the permissive approach for building the pseudo-reactions is to allow any of the specific lipids to form the generic lipid class (Heavner *et al*, 2012). For instance, the following set of pseudo-reactions can be defined for PI:

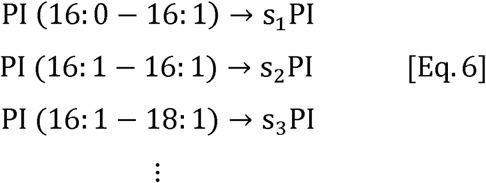

where s_i_ can be set to 1 or adapted to represent the cost of producing each specific lipid. The problem of this approach is that it disregards the acyl chain distribution, even if FAME data is available. Therefore, once simulations are computed, the model will always end up preferring the “cheapest” species to produce in terms of carbon and energy, usually corresponding to species with the shortest acyl chains, unless the s_i_ coefficients are arbitrarily tuned to favor longer chains.

Finally, even though for some specific species such as ergosterol the measured abundance [mg/gDW] can be directly transformed to the stoichiometric coefficient in Eq.4 [mmol/gDW], for the case of most lipids the measured abundance cannot be directly converted, as the molecular weight varies between specific lipid species. Hence, average molecular weights need to be estimated in both permissive and restrictive approaches, leading to skewed predictions.

### 2.2. Representing lipid constraints with the aid of SLIME reactions

In this study we solve the problems presented above through two new types of pseudo-reactions, to account for both constraints on lipid classes and on acyl chains. The first pseudo-reactions **S**plit **L**ipids **I**nto **M**easurable **E**ntities and are hence referred to as SLIME reactions. As the name suggests, these pseudo-reactions take each specific lipid and split it into its basic components, i.e. its backbone and acyl chains:

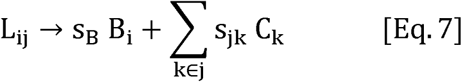

where L_ij_ is a lipid of class i and chain configuration j, B_i_ the corresponding backbone, C_k_ the corresponding chain k, and s_B_ and s_jk_ the associated stoichiometry coefficients. These reactions replace any pseudo-reaction of the sort of Eq.5 or Eq.6 that were already present in the model (Figure 1).

The second type of pseudo-reactions are new lipid pseudo-reactions, which will in turn replace Eq.4, the old lipid pseudo-reaction that only constrained lipid classes. There are now three different lipid pseudo reactions (Figure 1): the first pulls all backbone species created in Eq.7 into a generic backbone and uses the corresponding abundance data [g/gDW] as stoichiometric coefficients. The second reaction does the same with the specific acyl chains, with the data from FAME analysis [g/gDW], to create a generic acyl chain. Finally, the third reaction merges back together the generic backbone and the generic acyl chain into a generic lipid, which will be used in the biomass pseudo-reaction as in Eq.2.

For the new reactions to be consistent, we need to choose adequate stoichiometric coefficients for Eq.7. If the abundance data would be molar, s_B_ would be equal to 1 and s_jk_ would be equal to the number of repetitions of the corresponding acyl chain k in lipid j. However, as the abundance data often comes in mass units, s_B_ must be equal to the molecular weight [g/mmol] of the full lipid, and s_jk_ must be equal to the molecular weight of the corresponding acyl chain k, multiplied by the number of repetitions of k in configuration j. By choosing these values we allow the SLIME reactions to convert the molar production of the lipid [mmol/gDWh] into a mass basis [g/gDWh], which in turn will be converted to a lipid turnover [1/h] by the lipid pseudo reactions.

### 2.3. Data used

All data used in this study was collected from literature. For the main model analysis (section 3.1) and the analysis of lipid metabolism under increasing levels of stress (section 3.3), aerobic glucose-limited chemostat data of *S. cerevisiae*, strain CEN.PK113-7D, growing on minimal media at a growth rate of D = 0.1 h^-1^ was used (Lahtvee *et al*, 2017). The mentioned study collected lipid abundance data in mg/gDW for both lipid classes and acyl chains for 1 reference condition plus 9 different conditions of stress (temperature, ethanol and osmotic stress). Additionally, carbohydrate, protein and RNA content [g/gDW] was measured for all stress conditions, together with flux data [mmol/gDWh] for glucose and oxygen uptake, and glycerol, acetate, ethanol, pyruvate, succinate and CO_2_ production.

For model validation (section 3.2), aerobic data of *S. cerevisiae*, strain BY4741, grown on SD media at maximum growth rate (shake flask cultures) was used (Ejsing *et al*, 2009). The authors from that study introduced a novel quantification method for detecting the abundance of up to 250 singular species of lipids. Out of those, 102 were used in our study as they had direct correspondence to a species in the GEM employed. Those species represented on average 84% of the total detected lipid abundance, which was considered high enough to proceed with the rest of the analysis. The values were converted from mol/mol to mg/gDW assuming an 8% lipid abundance in biomass (Lahtvee *et al*, 2017) and considering the unmatched lipid percentage previously mentioned. Additionally, a protein composition of 0.5 g/gDW, an RNA composition of 0.06 g/gDW, a glucose uptake of 20.4 mmol/gDWh, and biomass growth rate of 0.41 h^-1^ were assumed based on previous batch simulations of the yeast GEM (Sánchez *et al*, 2017).

### 2.4. Model enhancement details

The consensus genome-scale model of yeast (version 7.8.0) was obtained from https://github.com/SysBioChalmers/yeast-GEM. Compared to the published version (Aung *et al*, 2013), the model included several manual curations detailed in previous work (Sánchez *et al*, 2017), a clustered biomass pseudo-reaction, and metabolite formulas added to every lipid. Combining this model together with the experimental data, the following 5 steps were followed to create a model with SLIME reactions, specific to each experimental condition:

1. Add pseudo-metabolites representing each specific backbone, each specific acyl chain, the generic backbone and the generic acyl chain.
2. Add for each specific lipid species a SLIME reaction, as described in section 2.2. These reactions replace the previous ones of the sort of Eq.6 in the model (Heavner *et al*, 2012).
3. Add all three lipid pseudo-reactions described in section 2.2 using the experimental data [g/gDW]. These reactions replace the original lipid pseudo-reaction.
4. Scale either the lipid class or the acyl chain abundance data so that they are proportional, as the approach is based on exact mass balances. For this, an optimization problem is carried out where the coefficients of the corresponding pseudo-reaction are rescaled to minimize to zero the excretion of unused backbones and acyl chains (Figure S1).
5. Finally, scale any other component in the biomass pseudo-reaction for which there is data, and ensure that the biomass composition adds up to 1 g/gDW (Chan *et al*, 2017) by rescaling the total amount of carbohydrates, which was not measured in the datasets employed.

To compare the performance of the new enhanced model, an additional model for each condition was created, which did not have the acyl chain pseudo-reaction, but instead exchange reactions for each acyl chain, so that the model could freely choose the acyl chain distribution. As this is equivalent to the permissive approach mentioned in section 2.1, we refer to this model in the following as the “permissive” model, and use it to benchmark our analysis. It should be mentioned here that experimental data showed that the acyl-chain distribution in yeast varies considerably across lipid classes (Figure S2). The “restrictive” approach assumes this as constant, which is a clear oversimplification and leads to skewed predictions; we therefore excluded this approach from our comparison.

### 2.5. Simulation details

For all FBA simulations, measured exchange fluxes were used to constrain the model, allowing up to a 5% of deviation from the average measurements, and a parsimonious FBA approach (Lewis *et al*, 2010) was followed, maximizing first the ATP turnover and then minimizing the total sum of absolute fluxes, in order to find the most compact solution. The obtained ATP turnover value is equal to the sum of the growth associated ATP maintenance (GAM) and the non-growth counterpart (NGAM, equal to 0.7 mmol/gDWh in the original model), and it was stored to compare ATP costs from transitioning from one state to another.

The variability of each different lipid species was computed in section 3.1 with flux variability analysis (FVA) (Mahadevan & Schilling, 2003) to each corresponding group of SLIME reactions at a time; e.g., for assessing the variability of C18:0 in PI, FVA was applied to all SLIME reactions producing PI and any C18:0 acyl chains. Variability was also assessed in section 3.2 with an artificial centering hit-and-run implementation of random sampling (Megchelenbrink *et al*, 2014). Abundances in mg/gDW of each lipid species were then computed from the corresponding SLIME reaction fluxes, multiplied by the molecular weight and divided by the biomass growth rate. Simulations were all performed using the COBRA toolbox for Matlab^®^ (Heirendt *et al*, 2017) with Gurobi^®^ 7.5 set as optimizer.

## 3. Results

### 3.1. Improved model of yeast

We implemented SLIMEr in the consensus genome-scale model of yeast version 7.8.0, which had at the start 2224 metabolites and 3496 reactions. Out of those reactions, 176 corresponded to reactions of the sort of Eq.6, which were replaced by 186 SLIME reactions that cover in total 19 lipid classes and 6 different acyl chains. An additional 27 metabolites (including both specific and generic backbones and acyl chains) and 15 reactions (including transport reactions, lipid pseudo-reactions and exchange reactions) were added to the model, and 10 metabolites and 1 reaction (connected to previously deleted reactions) were removed. The final enhanced model had therefore 2241 metabolites and 3520 reactions.

For the reference model, we used both lipid profiling and FAME data at low growth rate and no stress conditions (Lahtvee *et al*, 2017). The lipid profile was rescaled to be proportional to the FAME data, as detailed in section 2.4. After using SLIMEr, as expected, the enhanced model showed the same acyl chain distribution as the experimental data (Figure 2A), whereas the permissive model predicted mostly acyl chains of 16-carbon length (less costly), and only a small amount of 18-carbon length to satisfy the requirement of ergosterol ester (Figure S3), as ergosteryl oleate is cheaper to produce mass-wise than ergosteryl palmitoleate (Table S1).

**Figure 2:**
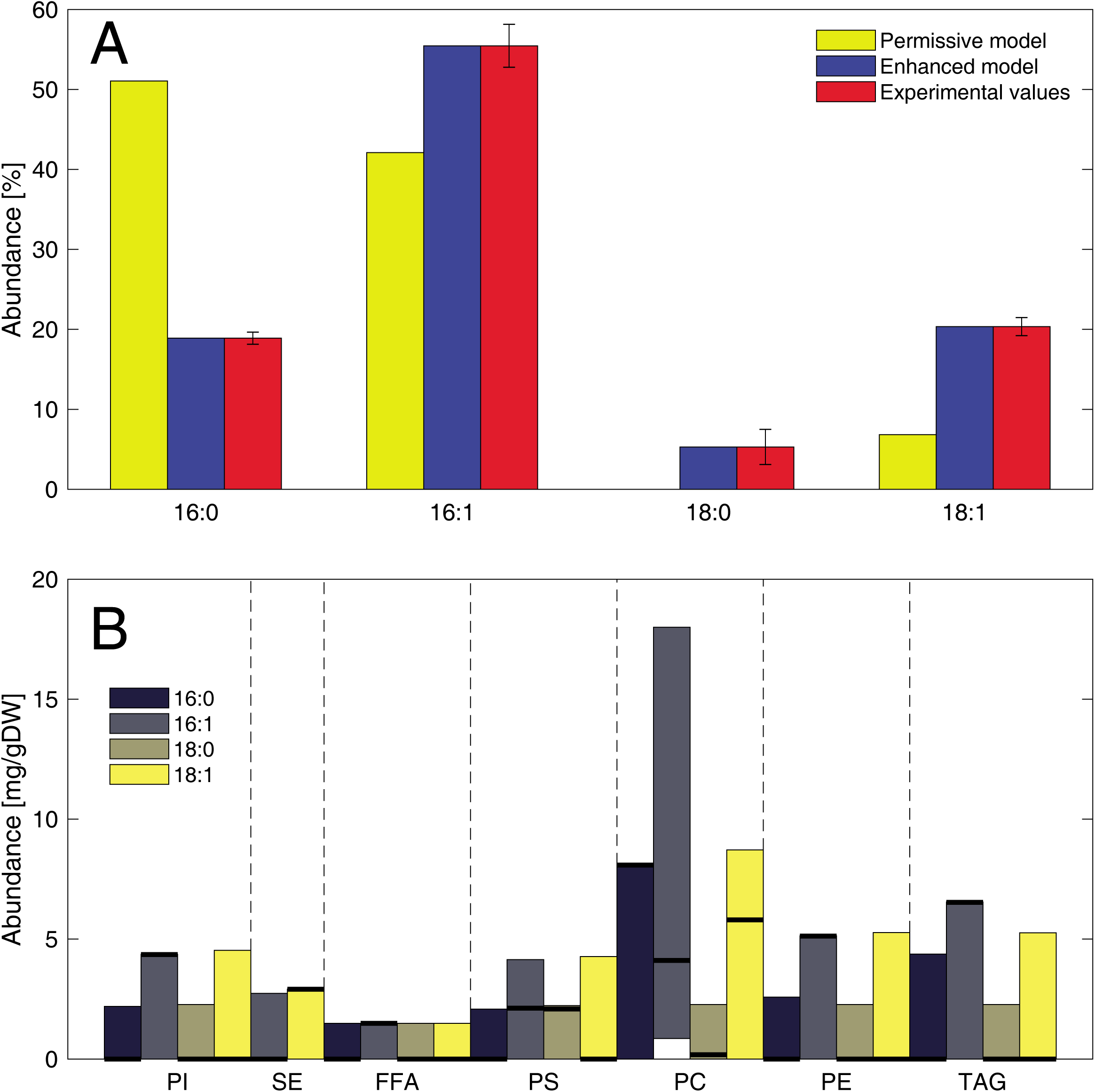
The enhanced GEM with improved constraints on lipid metabolism. **(A)** By using SLIMEr, a correct acyl chain composition is enforced. **(B)** Breakdown of the acyl chain distribution and variability predicted by the enhanced GEM, for each experimentally detected lipid class. Thick dark lines correspond to parsimonious FBA predictions, while the FVA allowed ranges are shown with colored bars.

With the enhanced model we also studied in how many ways lipid requirements can be satisfied spending the same amount of energy, by performing FVA (Figure 2B). Comparing these predictions to the ones of the permissive model (Figure S3), we saw some reductions in variability, coming mostly from changes in phosphatidylcholine and triglyceride content. However, despite the additional constraints imposed, lipid metabolism could still rearrange itself in a wide amount of combinations, and overall flux variability did not decrease significantly (Figure S4A). This agrees with experimental observations that lipid metabolism is highly flexible (Ejsing *et al*, 2009); therefore, handling lipid metabolism with SLIME reactions is preferred over alternative approaches, such as models that constrain single individual lipid species (Mardinoglu *et al*, 2014; Monk *et al*, 2017), as these limit the organism to only one feasible state of lipid metabolism and hence bias results.

### 3.2. Model predictions versus experimental data

To validate model predictions, we used data from literature (Ejsing *et al*, 2009) where each specific lipid species was measured (section 2.3). This data was added up to compute the totals of each lipid class and each acyl chain, and these sums were in turn used as input for creating both a permissive and an enhanced model. In this case, as a total lipid abundance of 8% was assumed, the acyl chains abundances were rescaled to be proportional to the lipid classes abundances (section 2.4).

We then performed random sampling on the resulting models, to generate 10,000 flux distributions for each model and for each of the 8 conditions of the study (4 different strains cultivated at 2 different temperatures). Comparing the *in silico* lipid compositions to the original *in vivo* measurements (Figure 3A), the enhanced model improved the average prediction error for all experimental conditions (Table S2), and overall the simulated lipid distributions came much closer to the experimental values compared to the permissive model (Figure 3B).

**Figure 3:**
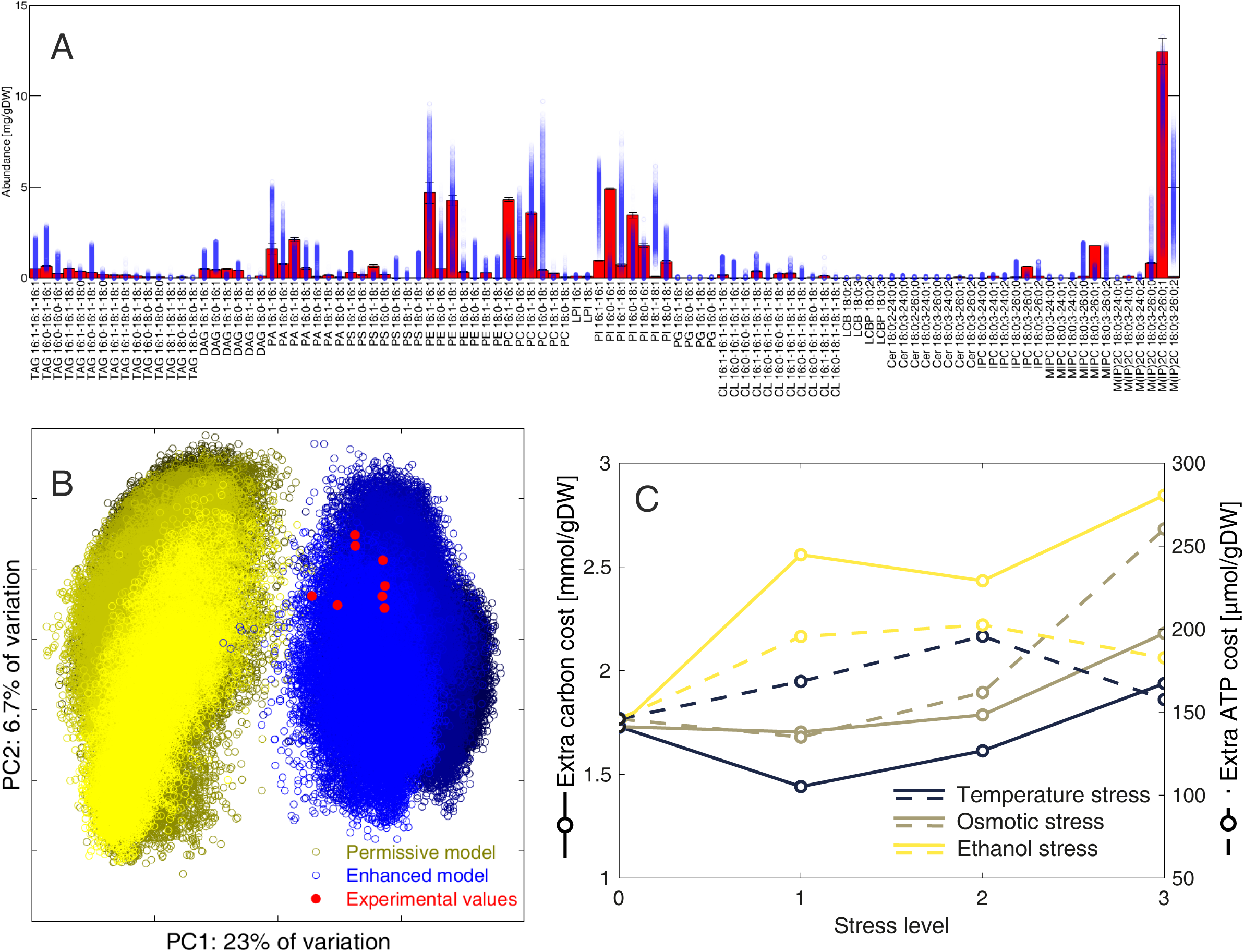
Using experimental data to validate the model and predict energy costs. **(A)** The lipid composition of 10,000 simulations of the enhanced model achieved with random sampling are presented for each specific lipid species (blue circles), compared to the actual measured experimental values (red bars). **(B)** Principal component analysis of all (log transformed) lipid abundance distributions for both the permissive (yellow) and enhanced (blue) models, compared to experimental values (red). Different tonalities of yellow and blue indicate the 8 different simulated conditions and strains. **(C)** Carbon costs (continuous lines) and ATP costs (segmented lines) of satisfying the acyl chain distribution at 3 increasing levels of stress of temperature (33, 36 and 38°C), NaCl concentrations (200, 400 and 600 mM) and EtOH concentrations (20, 40 and 60 g/L).

It should be noted that even though SLIMEr improved the model’s lipid composition predictions, many other distributions were still predicted to be equally likely, both for the reference condition (Figure 3A) and all strains (Figure 3B, Figure S5). Furthermore, the standard deviation remained overall constant compared to the “permissive” approach (Figure S4B, Figure S4C). This highlights the high level of regulation in place to adapt the distribution of lipid species in *S. cerevisiae* depending on the environmental conditions.

### 3.3. Energy costs at increasing levels of stress

By using SLIMEr, we can better represent the lipid expenses of transitioning from a metabolic state of low energy demand to high energy demand. To show this, we integrated data of yeast grown under different levels of stress (Lahtvee *et al*, 2017), as these stress levels were previously shown to be associated to an increase in maintenance energy (Lahtvee *et al*, 2016). Comparing simulations for the reference condition (same as in section 3.1), the permissive model could produce 145.9 μmol(ATP)/gDW more than the enhanced model, which corresponds to the extra energy required to achieve the acyl chain distribution. By repeating the analysis for all 9 conditions of stress, we found that both the extra energy and carbon expenses showed an increasing trend as the stress levels increased (Figure 3C), and so did the variability of flux distributions (Figure S6).

In the reference condition, the simulated growth-associated ATP maintenance (GAM) without accounting for known polymerization costs of proteins, carbohydrates, RNA and DNA (Förster *et al*, 2003) was 36.96 mmol(ATP)/gDW, which corresponds to the maintenance costs of unspecified functions in the model, such as protein turnover, maintenance of membrane potentials, etc. The ATP cost for achieving correct acyl chain distribution under reference conditions corresponded then to 0.4% of the total costs of processes not included in the model.

## 4. Discussion

With SLIMEr we can now correctly represent the biomass requirements from lipid metabolism in genome-scale metabolic models. The approach allows the model to satisfy at the same time requirements on the lipid class and acyl chain distributions, which is a significant improvement compared to only being able to constrain lipid classes (Aung *et al*, 2013; Brunk *et al*, 2018). We have also shown the high degree of flexibility in lipid metabolism, which shows that approaches that over-constrain the lipid requirements by enforcing specific concentrations for individual species (Mardinoglu *et al*, 2014; Monk *et al*, 2017; Lachance *et al*, 2018) or forcing a given acyl chain distribution to all species (Nookaew *et al*, 2008; Kerkhoven *et al*, 2016) are not suitable for handling this flexibility. Finally, we have demonstrated the use of the expanded model as a tool to compute lipid requirements in varying experimental conditions. We expect the enhanced model to be very useful for metabolic engineering applications, particularly for designing strains that can rearrange the chain length distribution of specific lipid classes (Bergenholm *et al*, 2018).

As previously mentioned, we did not see a significant reduction in flux variability of predictions (Figure S4). This is partly explained as in each simulation we maximize the ATP maintenance; therefore, simulations of the permissive model (which did not have constraints on the acyl chain distribution) had a slightly higher ATP maintenance, making simulations overall similarly constrained. However, the main advantage of using SLIMEr is not to constrain simulations more, but instead to constrain lipid fluxes such that they better match biologically feasible distributions (Figure 3B). It is also important to note that the model does not take other physiological properties into account, such as specific regulation, or curvature and fluidity of membranes as function of lipid composition and/or temperature. It only takes FAME analysis and lipid profile data, and demonstrates that specific lipid distributions are consistent with these measurements. It would be of interest to account for additional data and processes such as the ones mentioned, but this is beyond the scope of this study.

Even though developed for the consensus GEM of *S. cerevisiae*, this approach can be extended to any other model and/or organism. The main challenge here is to map all lipids in the model to the corresponding pseudo-metabolites (backbones and chains), as conventions for naming lipids vary a great deal between different databases and models. Introduction of standardized metabolite ids (Dräger & Palsson, 2014; Moretti *et al*, 2016) can significantly aid this otherwise laborious task.

## Declarations

### Ethics approval and consent to participate

Not applicable.

### Consent for publication

Not applicable.

### Availability of data and material

All data analyzed in this study are included in the following 2 published articles [and their supplementary information files]:

- Lahtvee PJ, Sánchez BJ, Smialowska A, Kasvandik S, Elsemman IE, Gatto F & Nielsen J (2017) Absolute Quantification of Protein and mRNA Abundances Demonstrate Variability in Gene-Specific Translation Efficiency in Yeast. *Cell Syst.* **4:** 495–504.e5
- Ejsing CS, Sampaio JL, Surendranath V, Duchoslav E, Ekroos K, Klemm RW, Simons K & Shevchenko A (2009) Global analysis of the yeast lipidome by quantitative shotgun mass spectrometry. *Proc. Natl. Acad. Sci.* **106:** 2136–2141

SLIMEr, together with all scripts necessary to reproduce the results presented in this study, are available at https://github.com/SysBioChalmers/SLIMEr. All new SLIME reactions and lipid pseudo-reactions have been added to the consensus GEM of yeast and are available in version 8.1.0: https://github.com/SysBioChalmers/yeast-GEM/releases/tag/v8.1.0.

### Authors’ contributions

JN and BJS conceived the project. BJS, FL and EJK designed the mathematical formulation. BJS implemented the algorithm and performed all computational simulations. BJS and FL processed the literature data. BJS wrote the original draft. All authors read, edited and approved the final manuscript.

## Acknowledgements

The authors would like to thank Dr. Hongzhong Lu for help in annotation of lipid formulas, Dr. Petri-Jaan Lahtvee and Dr. Paulo Teixeira for aiding in data analysis, and Sebastián Mendoza for guidance with the random sampling analysis.

## Abbreviations

CL: Cardiolipin
Cer: Ceramide
COBRA: Constraint-based reconstruction and analysis
DAG: Diglyceride
FAME: Fatty acid methyl esters
FBA: Flux balance analysis
FVA: Flux variability analysis
GAM: Growth associated ATP maintenance
GEM: Genome-scale metabolic model
IPC: Inositolphosphoceramide
LCB: Long-chain base
LCBP: Long-chain base phosphate
LPI: Lysophosphatidylinositol
M(IP)2C: Mannosyl-diinositolphosphoceramide
MIPC: Mannosyl-inositolphosphoceramide
NGAM: Non-growth associated ATP maintenance
PA: Phosphatidate
PC: Phosphatidylcholine
PE: Phosphatidylethanolamine
PG: Phosphatidylglycerol
PI: Phosphatidylinositol
PS: Phosphatidylserine
SLIME: Split lipid into measurable entities
TAG: Triglyceride

